# Matrix selection for the visualization of small molecules and lipids in brain tumors using untargeted MALDI-TOF mass spectrometry imaging

**DOI:** 10.1101/2023.09.25.559427

**Authors:** Tianyao Lu, Lutz Freytag, Vinod K. Narayana, Zachery Moore, Shannon J. Oliver, Adam Valkovic, Brunda Nijagal, Amanda Peterson, David P. de Souza, Malcolm J. McConville, James R. Whittle, Sarah A. Best, Saskia Freytag

## Abstract

Matrix-assisted laser desorption/ionization mass spectrometry imaging allows the study of metabolic activity in the tumor microenvironment of brain cancers. The detectable metabolites within these tumors are contingent upon the choice of matrix, deposition technique, and polarity setting. In this study, we compared the performance of three different matrices, two deposition techniques, and use of positive and negative polarity in two different brain cancer types and across two species. Optimal combinations were confirmed by comparative analysis of lipid and small molecule abundance using liquid chromatography–mass spectrometry and RNA-sequencing assessing differential metabolites between normal and tumor regions. Our findings indicate that the recrystallized α cyano-4-hydroxycinnamic acid matrix in positive polarity offered superior performance for both detected metabolites and consistency with other techniques. Beyond these implications for brain cancer, our work establishes a workflow to identify optimal matrices for spatial metabolomics studies.

## 1. Introduction

Brain cancer is a devastating malignancy that is currently incurable, with significant adverse effects on the patient’s quality of life. Like many other cancers, tumor cells in brain cancer display an altered metabolism, with an increase in the tricarboxylic acid (TCA) cycle related to tumor progression [1]. One of the metabolic pathways that is commonly altered is the switch to aerobic glycolysis, also known as the Warburg effect [2]. This metabolic adaptation, which is typically observed in healthy proliferating cancer cells, allows the cells to produce energy more efficiently resulting in rapid cell growth and division. Other metabolic alterations include the production of the onco-metabolite 2-hydroxyglutarate (2-HG) as a result of a point mutation in the isocitrate dehydrogenase 1/2 (*IDH1/2*) gene, strongly associated with a discrete subset of brain cancers (*IDH*-mutant gliomas) [3–5]. 2-HG inhibits the activity of enzymes that are involved in DNA and histone modifications, leading to epigenetic changes in gene expression that promote cancer cell growth and survival.

Metabolic activity is not uniformly altered in brain cancer tumors, reflecting complex and multifactorial regulation, that in turn depends on the cellular microenvironment and underlying genetic profile of the malignant cells [6,7]. For example, oxygen availability determines the rate of aerobic glycolysis utilization throughout brain cancer tumors. Matrix-Assisted Laser Desorption/Ionization (MALDI) Mass Spectrometry Imaging (MSI) is a robust tool for studying the spatial distribution of metabolic alterations [8]. MALDI MSI rasterizes thin sections of tissue to produce mass spectra for each individual pixel allowing the detection of various analytes, such as small molecules, lipids and peptides. Thus, MALDI MSI enables highly sensitive analysis with a spatial resolution ranging between 10-200 μm. Due to its favorable balance between ease of sample preparation, chemical specificity/sensitivity, and spatial resolution, MALDI MSI has become the most widely used tool for spatial metabolomics in recent years.

MALDI MSI can use different analyzers, including: Time-Of-Flight (TOF), Fourier transform ion cyclotron resonance, Orbitrap, and quadrupole ion trap, among which the former are the most common. The TOF analyzer measures the time it takes for ions to travel a certain distance, which is proportional to their mass-to-charge (m/z) ratio, while the Fourier transform ion cyclotron resonance uses a magnetic field to separate ions based on their m/z ratio. While the latter provides favorable mass resolution and mass accuracy for unambiguous identification of molecules, MALDI TOF MSI provides considerably faster acquisition time (several hours for tissue sections with an area of ∼1 cm^2^) and broader mass range [9]. Hence, MALDI TOF MSI has become the prevalent technique for investigating the spatial distribution of analytes in biological samples, making it a valuable tool in the field of biomarker discovery and disease diagnosis [10].

The selection of a suitable matrix, which is used to assist in the desorption and ionization of molecules from the sample surface, is crucial for the success of MALDI TOF MSI [11]. There are many matrices each suitable for different analytes of interest. Common considerations include i) optical absorption, ii) interference with analytes of interest, iii) polarity setting, iv) stability, v) sensitivity, vi) vaporization ease, and vii) mass range of the analytes of interest [12]. Despite considerable efforts dedicated to developing a universally optimal matrix, currently matrix selection still needs to be optimized for each experiment. Here, we have compared five different matrices using both positive and negative polarities in MALDI TOF MSI for the detection of small molecules and lipids in murine and human brain tumors. We not only establish recommendations for matrix selection for this experimental setting but present a strategy, including robust and automated computational pipelines, for quantitative assessment guiding matrix selection applicable to any experimental setting.

## 2. Materials and Methods

### 2.1. Mouse Model

All experiments presented in this study were conducted according to regulatory standards approved by the Walter and Eliza Hall Institute Animal Ethics Committee. The CT-2A syngeneic mouse cell line (10,000 cells) [13] was transplanted into the cortex of female 7 week old C57Bl/6 mice according to standard procedures using the coordinates 1 mm lateral and 1 mm anterior to the bregma to a depth of 2.5 mm [14]. Mice were monitored for neurological disturbances and collected as a cohort at ethical endpoint. Tumors were harvested within the intact brains, transferred to the laboratory in PBS and immediately flash frozen directly in isopentane in a liquid nitrogen bath. Brains were stored at -80 °C until downstream sample preparation. Where relevant for bulk studies, tumor and normal regions were dissected directly from the frozen material on dry ice and processed independently.

### 2.2. Human Samples

Patient samples were obtained directly from surgery via the Royal Melbourne Hospital Neurosurgery Brain and Spine Tumor Tissue Bank (Melbourne Health Ethics #2020.214) and immediately flash frozen directly in isopentane in a liquid nitrogen bath. Patient tumors were stored at -80 °C until downstream sample preparation.

### 2.3. Sample Preparation

#### 2.3.1. MALDI TOF-Based Mass Spectrometry Experiment

Frozen tissue was sectioned at a thickness of 10 µm directly onto Indium Tin Oxide (ITO) coated glass slides (surface resistance of 70-100 Ω). Optimal cutting temperature (OCT) reagent was used to mount the tissue onto the cryotome chuck (-17 °C) for sectioning and was not exposed in the region of sectioning for metabolomics analysis due to interference. Sections were stored at -80 °C until analysis and dried in a vacuum desiccator for 30 min, followed by collection of optical images using the light microscope embedded in the MALDI TOF MSI instrument (iMScope QT) prior to matrix application.

To compare the different matrices and application methods (Table 1, Supplementary Material Figure 1a), serial sections on separate ITO glass slides were coated with the various MALDI matrices; MALDI grade 9-Aminoacridine (9-AA, P# 92817), 2,5-Dihydroxybenzoic acid (DHB, P# 149357), α-cyano-4-hydroxycinnamic acid (CHCA, P# C2020) purchased from Sigma-Aldrich, Germany. Matrix deposition was performed either by single step matrix vapor deposition using iMLayer (Shimadzu, Japan) to sublimate and deposit an even layer of small crystals across the surface of the tissue, or by 2 step deposition method using iMLayer for sublimation and iMLayer AERO (Shimadzu, Japan) for matrix spraying to recrystallize and obtain fine matrix crystals, that enable high sensitivity and high spatial resolution (Supplementary Material Figure 1b). The thickness of the vapor-deposited matrix was 1.5, 0.9 and 0.7 µm, and the deposition temperature was 180, 220 and 250 °C, for DHB, 9-AA and CHCA matrices respectively. To recrystallize the matrix vapor-deposited tissue samples, 9-AA and CHCA matrix solutions were used for spraying through iMLayer AERO. Both thickness and temperature, remained the same for each of the matrices regardless of recrystallization. 9-AA matrix concentration was kept at 10 mg/mL in methanol/water (90:10, v/v) solution. Four layers of 9-AA were sprayed at a 90 mm/sec stage speed, with 1 sec dry time at a 5 cm nozzle distance. The pumping pressure and the spraying pressure were kept constant at 0.05 and 0.15 MPa. For CHCA matrix spraying, 8 layers of 10 mg/mL CHCA in acetonitrile/water (50:50, v/v) with 0.1% trifluoroacetic acid solution were used. The stage was kept at 70 mm/sec with 1 sec dry time at a 5 cm nozzle distance and pumping pressure kept constant at 0.1 and 0.2 MPa respectively.

**Table 1.** Summary of matrices, deposition methods, adduct ion, and ranges.

| Matrix | Abbreviation | Deposition Method | Adduct ion/ polarity | Ranges (m/z) |
| --- | --- | --- | --- | --- |
| 9-Aminoacridine | 9-AA- | Sublimation only | Anionised ([M-H]-) | 70-400<br>400-1,000 |
| 9-Aminoacridine | 9-AA (2step)- | Sublimation followed by recrystallisation | Anionised ([M-H]-) | 70-400<br>400-1,000 |
| 9-Aminoacridine | 9-AA+ | Sublimation only | Cationised ([M+Na]+; [M+K]+; [M+H]+) | 70-400 |
| 9-Aminoacridine | 9-AA (2step)+ | Sublimation followed by recrystallisation | Cationised ([M+Na]+; [M+K]+; [M+H]+) | 70-400 |
| α-cyano-4-hydroxycinnamic acid | CHCA- | Sublimation only | Anionised ([M-H]-) | 70-400 |
| α-cyano-4-hydroxycinnamic acid | CHCA (2step)- | Sublimation followed by recrystallisation | Anionised ([M-H]-) | 70-400 |
| α-cyano-4-hydroxycinnamic acid | CHCA+ | Sublimation only | Cationised ([M+Na]+; [M+K]+; [M+H]+) | 70-400<br>400-1,000 |
| α-cyano-4-hydroxycinnamic acid | CHCA (2step)+ | Sublimation followed by recrystallisation | Cationised ([M+Na]+; [M+K]+; [M+H]+) | 70-400<br>400-1,000 |
| 2,5- dihydroxy benzoic acid | DHB- | Sublimation only | Anionised ([M-H]-) | 70-400 |
| 2,5- dihydroxy benzoic acid | DHB+ | Sublimation only | Cationised ([M+Na]+; [M+K]+; [M+H]+) | 70-400 |

All MSI experiments were performed using iMScope QT instrument (Shimadzu, Japan). The instrument is equipped with a Laser-diode-excited Nd:YAG laser, and an atmospheric pressure MALDI. To achieve better MS sensitivity the data of two neighboring m/z ranges 70-400 for polar metabolites and m/z 400-1000 Da for lipids with positive and negative polarity were acquired. Laser diameter (spatial resolution) was kept at 10 µm and same pitch (variable stage step down) was matched to the laser diameter. For all the samples data was acquired using a laser intensity at 60 for 10 µm spatial resolution, detector voltage was set at 2.36 kV, laser repetition frequency set at 2000 Hz, desolvation line temperature maintained at 250 °C and a laser irradiation count of 50 shots were accumulated per pixel.

#### 2.3.2. Liquid chromatography–mass spectrometry

Pre-weighed tissue samples of 20 mg each were extracted for Liquid Chromatography–Mass Spectrometry (LC-MS) analysis by homogenisation in a Precellys^®^ 24 Tissue homogeniser coupled to a Cryolys^®^ cooling system (Bertin Technologies) in 500 μL of 3:1 Methanol:Milli-Q water (containing internal standards). Extracts were vortexed and mixed on a thermomixer (10 mins) to ensure complete metabolite extraction and then centrifuged at 4 °C for 10 minutes at 12,700 rpm to remove the tissue pellet.

For polar metabolite analysis: 80 μL of the methanol/water supernatant was transferred into a glass HPLC vial containing a glass insert. A further 20 μL from each sample was pooled to generate a pooled biological quality control (pbQC) samples, which was run after every five biological samples.

Analyses of polar analytes in the samples was performed on the Orbitrap ID-X Tribrid mass spectrometer (Thermo Scientific) coupled to a Vanquish Horizon UHPLC system (Thermo Scientific). Separation of polar metabolites was performed on a Merck SeQuant ZIC-HILIC column (150 mm × 4.6 mm, 5 μm particle size) maintained at 25 °C, using a binary gradient consisting of solvent A: 20 mM ammonium carbonate (pH 9.0; Sigma–Aldrich) and solvent B: 100 % ACN. The gradient run was as follows: time (t) = 0.0 min, 80 % B; t = 0.5 min, 80 % B; t = 15.5 min, 50 % B; t = 17.5 min, 30 % B; t = 18.5 min, 5 %; t = 21.0 min, 5 % B; t = 23–33 min, 80 % at a solvent flow rate of 300 μl/min.

Data analysis was undertaken in El Maven software (v0.12.1). Level 1 metabolite identification, according to the Metabolite Standard Initiative [15], was based on matching accurate mass and retention time to the 550 authentic standards in the Metabolomics Australia in-house library.

For targeted lipid analysis: 600 μL of 100 % chloroform (containing lipid internal standards) was added to the remaining homogenate to bring the ratio to 2:1 chloroform:methanol. Extracts were vigorously vortexed to resuspend and further extract the lipids from the tissue pellet. The samples were then mixed on a thermomixer (10 mins at 10 °C) and centrifuged at 10 °C for 5 minutes at maximum speed to pellet the protein. The lipid extract was dried in 5 x 100 μL aliquots on a rotational vacuum concentrator (Christ RVC 2-33 CDplus) at 35 °C. Aliquots were dried, then reconstituted in 100 μL of 9:1 butanol:methanol. For a pbQC, 10 μL of each sample was pooled into a fresh vial.

Analyses of tissue lipids were performed on the Agilent 6490 LC-QQQ mass spectrometer coupled with an Agilent 1290 series HPLC system. Separation of lipids were conducted on a ZORBAX RRHD UHPLC C18 column (2.1×100 mm 1.8 mm, Agilent Technologies) maintained at 45 °C. Targeted mass spectrometry analysis was conducted in ESI positive ion mode with dynamic scheduled multiple reaction monitoring. Mass spectrometry settings and multiple reaction monitoring transitions for each lipid subclass and individual species was run as previously described [16]. Data analysis and integration of the chromatographic peaks was undertaken in MassHunter (B 10.1, Agilent Technologies) software.

#### 2.3.3. Bulk RNA sequencing

Extracted normal and tumor regions were ground in liquid nitrogen and RNA extracted using the RNeasy RNA extraction kit (Qiagen 74104) then analyzed using a 2200 Tapestation Analyser (Agilent). An input of 500 ng RNA was used to generate libraries (TruSeq RNA Library Prep v2, Illumina). Sequencing was performed on the NextSeq System (Illumina) to produce 132bp single-end reads.

### 2.4. Computational Analysis

#### 2.4.1. File conversion

MALDI TOF MSI data were obtained in the Shimadzu IMDX format. To facilitate subsequent analytical procedures, we converted the data into ImzML format using the “ImzML converter” (version 1.21.0 11302) functionality within the Shimadzu IMAGEREVEAL MS platform.

#### 2.4.2. Cardinal pipeline

We used the Bioconductor Cardinal package (version 3.0.2) with R-4.3.0, a specialized tool for the manipulation of MSI data, to process all MALDI TOF MSI data. Our pipeline involved the following steps (Figure 1a):

- Field of View (FOV) selection;
- Spectrum acquisition and preprocessing;
- Reference peak identification and refinement;
- Spectrum binning;
- Background peak removal.

**Figure 1.**
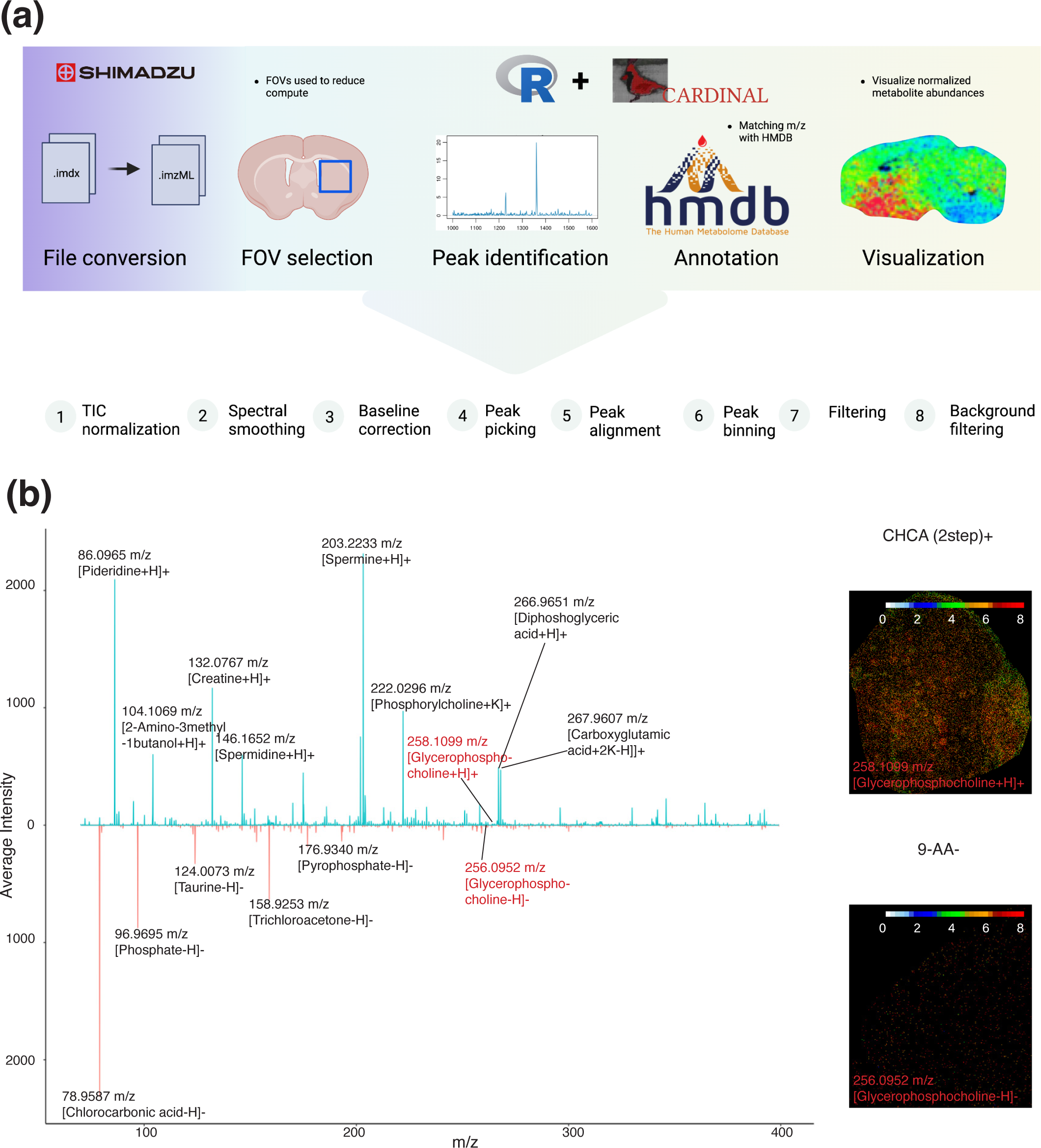
Computational pipeline (**a**) Computational pipeline for the analysis of MALDI-TOF data; (**b**) Comparison of spectra of small metabolite screen using the recrystallized CHCA matrix in positive polarity mode (top) and 9-AA in negative polarity mode (bottom) and spatial abundance of Glycerophosphocholine.

The initial spectral data were imported into the R environment as an “S4 object of class MSContinuousImagingExperiment” using the Cardinal::readImzML function [17]. To prevent memory-related issues during processing, a FOV of size 300 pixels by 300 pixels was selected. FOVs encompassed both the tumor region and the healthy brain region. Subsequently, normalization was performed utilizing the total ion current (TIC) method, followed by spectrum smoothing via Savitzky-Golay filter. Baseline distortions within the spectral data were addressed by applying median interpolation methods across 500 segments of the original spectrum.

To identify reference spectral peaks with significance, the acquired spectrum was processed through the Cardinal::peakpick function. Alignment of the spectrum was conducted using Cardinal::peakalign, incorporating a default estimated tolerance to yield candidate peaks. These candidate peaks were then subjected to a refinement process using the Cardinal::peakFilter function. This involved selecting local maxima across five spectral windows, while enforcing a minimum signal-to-noise ratio of 6. Additionally, a criterion was set requiring the signal to be detected in at least 1 % of all pixels. The processing of background peak bins followed the same procedures, but they were selected based on a minimum detection frequency in at least 5 % of the pixels. This criterion was applied because background signals tend to be less densely distributed compared to signals originating from the tissue.

Next, the processed spectrum was subjected to binning, employing the Cardinal::peakBin function on the filtered reference peaks. These peak bins encapsulated the final spectrum information. Finally, if an experimental peak is detected within 50 parts per million (ppm) at its nearest background peak, then the signal peak is removed to avoid noise. These peaks were subsequently utilized for downstream analyses and annotation tasks (Figure 1b).

#### 2.4.3. Annotation of peaks

The peaks identified by Cardinal were matched against the Human Metabolome Database (HMDB, version 5.0) [18] using the k-nearest neighbor methodology (Figure 1b). For each peak, a set of 30 nearest neighbors from generated HMDB peak list is identified. To refine the selection, metabolites with distances exceeding 50 ppm from the peak bin center are filtered out. The selection of 50 ppm as the filtering threshold is underpinned by the characteristics of the Shimadzu iMScope QT mass resolution. Meanwhile, our empirical investigation involving estimation around the peak produced by the matrix demonstrated that the average TOF-resolution was closer to 50 ppm. It is important to acknowledge that a single annotation of an HMDB entry can emanate from distinct peaks, attributable to different adduct ions.

#### 2.4.4. Pathway analysis

We employed overrepresentation-based pathway analysis via the stats::fisher.test (version 4.3.0) to assess the significance of associations between the metabolite hits and reference pathways obtained from the Relational Database of Metabolomics Pathways (RaMP, database version 2.3.0) [19]. The RaMP database captures both genetic and metabolomic information linked to specific pathways. The elements within these pathways are transformed into two distinct components: a metabolomics-specific portion and a gene-specific portion. These components were then stored as individual list items within the R programming environment. The p-value was adjusted by False Discovery Rate (FDR) using Benjamini Hochberg for multiple testing.

#### 2.4.5. Kernel density estimation

To effectively compare the distribution of peak bins between matrices, a comprehensive approach involving the estimation of the distribution of peak through the application of a Gaussian Kernel Density Estimation (KDE) was used. It is important to note that our objective here is distinct from the typical KDE approach, which aims to generate smooth curves. The computation of KDE was performed in R using the stats::density (version 4.3.0) function, with unbiased cross-validation bandwidth and parameter n=2500 (number of returned x-y coordinates) for small molecules and n=5000 for lipids.

#### 2.4.6. Wasserstein distance

The average intensity of a peak within a 50 pixel by 50 pixel Region Of Interest (ROI) centered at the pathologically distinct tumor injected region and a 50 pixel by 50 pixel ROI within the unaffected brain region were compared using a two-dimensional Wasserstein distance, the data were converted to an object of class “pp” with CRAN package transport (version 0.14.6), and the Wasserstein distance was calculated with transport::wasserstein function.

Additionally, the Wasserstein distance was used to compare between peak bins detected in different matrices. The comparison involved assessing the KDE of peak bin distribution across various matrices. Since the m/z ratio of the peaks represents diverse molecules in different ion mode, the m/z values of individual coordinates used to calculate the Wasserstein distance are corrected by subtracting the adduct ion mass. Given that each matrix exhibits a distribution of densities along the m/z detection range, the comparison was conducted within this context.

#### 2.4.6. Bulk RNA-sequencing processing

The bulk RNA-sequencing (RNA-seq) data was aligned using STAR with reference to mouse mm10 genome sequences. The aligned RNA-seq data were normalized by edgeR package (version 3.42.4) with edgeR::calcNormFactors and edgeR::cpm to obtain the normalized count [20].

#### 2.4.7. Comparison with LC-MS data

LC-MS metabolite data were pre-processed, including peak intensity normalization with the TIC method, background intensity subtraction, and annotation. Comparison of LC-MS data with MALDI TOF MSI data involved alignment of HMDB IDs, chemical formula, and synonyms. To compute the consistency between LC-MS and MALDI TOF MSI data, we computed the common regulatory trend in tumor versus the normal brain region. Specifically, in cases where a potential m/z value corresponds to the HMDB ID have a regulatory trend (upregulated or downregulated in tumor region) aligned with trends observed in the LC-MS data, metabolites associated with the HMDB ID are regarded as consistent. This strategic determination accommodates the intricate interplay of data sources and the diverse regulatory dynamics manifesting in the tumor region. Due to the absence of information about lipid backbones, we used a match of m/z with 2 and 10 ppm tolerance to align the LC-MS and MALDI TOF MSI lipid data.

#### 2.4.8. Comparison with RNA-sequencing data

To allow the comparison between bulk RNA-sequencing data and spatial metabolomics data, while reflecting the relative regulations of specific pathways in the tumor region, rank-based pathway enrichment was utilized. The rank-based pathway enrichment was conducted by fgsea::fgsea (version 1.24.0) function [21], the minimal size of sets of three was considered, while the maximal size was set to 1,500. The test was permuted for 2,500 times. Reference molecular signatures in different pathways were derived from RaMP.

## 3. Results

### 3.1. Matrix selection for small molecules in murine glioma

We compared the number of unique peaks detected for each matrix using MALDI TOF MSI in positive and negative polarity mode, across both tumor and normal tissue in mice with a transplanted CT-2A glioma. We detected 1,417 processed peaks using the CHCA 2 step recrystallization method in positive polarity mode followed by 801 and 709 peaks using CHCA and 9-AA sublimation in positive polarity mode, respectively (Figure 2a). The worst performing matrix was DHB, which only detected fewer than 140 peaks in both polarities. Unsurprisingly, we also found the least number of peaks attributed to the matrix for DHB in both polarities and the most background peaks for the recrystallized CHCA matrix in positive polarity mode (Supplementary Material Figure S2a). For all matrices and in either polarity, we identified a larger number of uniquely annotated chemical structures than unique peak bins with an average of 2.16 chemical formulas (isomers share the same formula) being matched to each peak. Meanwhile, on average 2.63 chemical formulas and 1.35 chemical formulas are matched on a single peak in positive and negative mode, respectively, owing to the relatively poor resolution of MALDI-TOF MSI.

**Figure 2.**
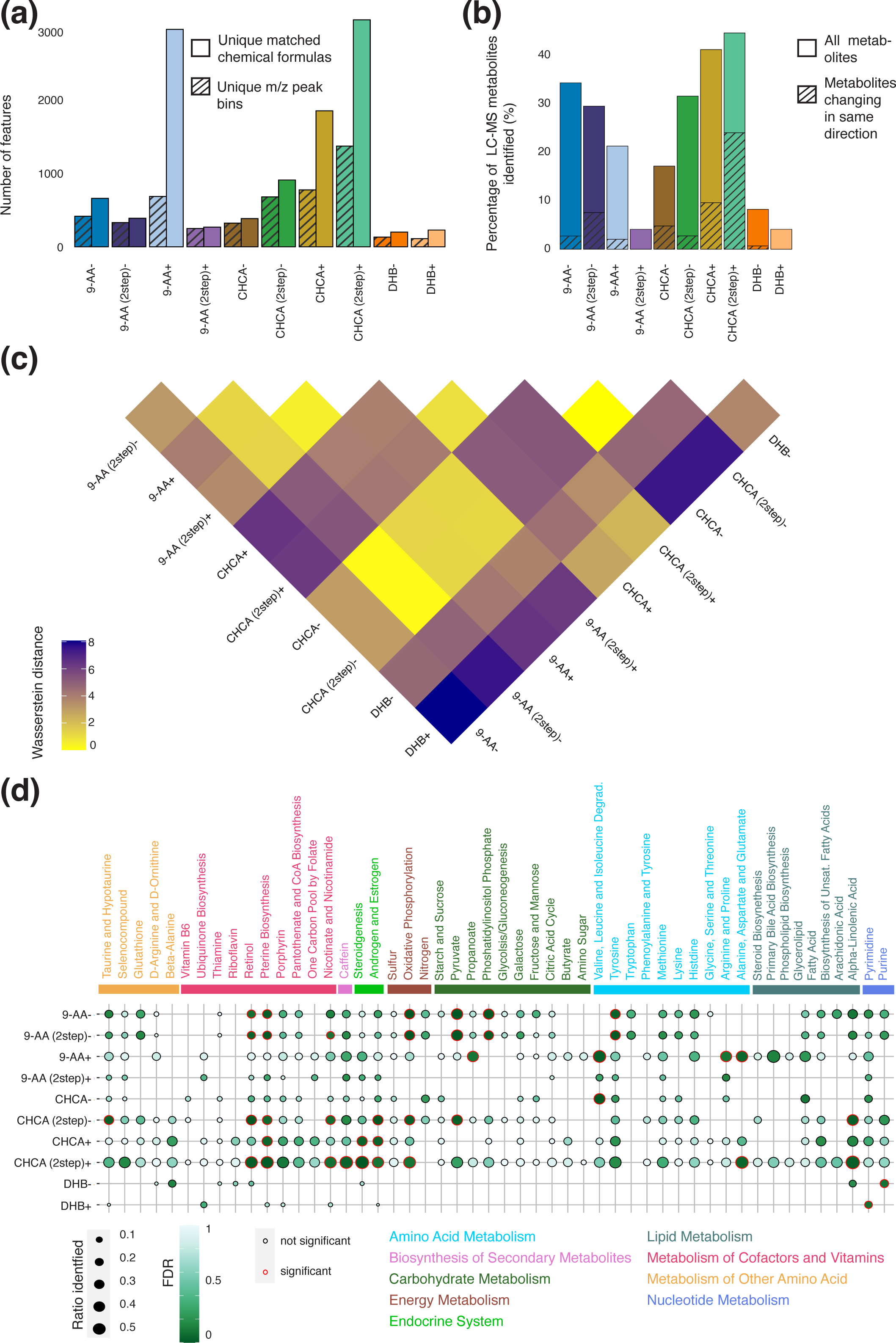
Comparison of different matrix and polarity combinations for detection of small molecules in murine glioma. (**a**) Number of unique identified annotated peaks and m/z peaks for each combination; (**b**) Percentage of metabolites detected by LC-MS identified by each combination, shaded area indicates the percentage of metabolites that had the same direction between the tumor and normal regions between LC-MS and MALDI-TOF; (**c**) Wasserstein distance between each combination with higher Wasserstein distances between two combinations indicate larger differences between identified peak distribution; (**d**) Ratio of detected metabolites per KEGG pathway and associated FDR for each combination, significantly enriched pathways are indicated with a red outline.

LC-MS on both tumor and normal samples from mouse with CT-2A glioma detected a total of 145 unique annotated small metabolites (70-400 m/z range). Comparing the detected peaks with MALDI-TOF MSI, we found that while the recrystallized CHCA matrix in positive polarity mode detected approximately 2-fold the number of raw peaks compared to CHCA sublimation only, both matrices matched approximately 40 % of the LC-MS metabolites. In contrast, the 9-AA in positive polarity mode, which had the second highest number of raw peaks detected, only identified 34 % of LC-MS metabolites. 9-AA using both deposition methods in positive polarity mode, along with DHB in both polarities, also identified few metabolites that had been present in previous studies of brain cancer (Supplementary Material Figure S2b). Recrystallized CHCA, in both polarities, as well as 9-AA using both deposition methods in negative polarity mode detected 13 of 15 metabolites of interest, detected in previous studies of brain cancer [22,23].

The 9-AA matrices using both polarities identified similar spectra (Figure 2c). These spectra were similar to those found by the CHCA matrices (using both deposition techniques) in negative polarity mode. CHCA matrices in positive polarity mode and DHB matrices in both polarities identified spectra that were more dissimilar to spectra identified by other matrices and polarity combinations and each other. It is therefore not surprising that peaks with annotatable entries in HMDB identified with the DHB matrix in negative polarity mode were largely not detected by other matrix and polarity combinations (Supplementary Material Figure S2c). In contrast, 71 % of the matched chemical formula identified by the recrystallized CHCA matrix in positive polarity mode were also found by other matrix and polarity combinations, possibly owing to the large total number of identified peaks. Interestingly, for 9-AA in positive polarity mode, 18 % of all matched chemical formulas were exclusively detected despite the low overall number of peaks detected.

To assess whether the different matrix and polarity combinations detected similar biological programs, we conducted an over-representation-based pathway analysis. In general, most matrix and polarity combinations, apart from those involving DHB, detected metabolites in the same KEGG pathways (Figure 2d). Purine metabolites, indicative of dividing cells and therefore crucial to examine cancer growth, were completely undetected in several matrix and polarity combinations, including 9-AA in positive polarity mode. Furthermore, only CHCA, using both deposition techniques, in positive polarity mode showed good coverage of metabolic pathways related to cofactors and vitamins. Pathway enrichment across multiple pathway databases indicated that recrystallized CHCA matrix in positive polarity mode was biased towards detecting metabolic pathways related to Hormone and Fatty Acid Mediated Insulin Secretion, Amino Acid Metabolism and Catabolism, Nucleotide Biosynthesis and Metabolism and Sulfur Amino Acid Metabolism (Supplementary Material Figure S2d). Interestingly, given the sample origin there was no bias for detecting metabolic pathways related to synaptic neurotransmission for this matrix and polarity combination.

### 3.2. Matrix selection for lipid detection in murine CT-2A glioma

To assess the utility of different matrices for lipid analysis, using only the 9-AA and CHCA matrices, we first compared the total number of unique peaks detected for each tested matrix and polarity combination (m/z screening range between 400-1000). We again detected the largest number of peaks 5,959 with the recrystallized CHCA matrix in positive polarity mode (Figure 3a). We also found the largest number of peaks attributed to this matrix and polarity combination (Supplementary Material Figure S3a). Out of all tested matrix and polarity combinations, 9-AA in negative polarity mode detected the least number of total and background peaks. For all matrices and in either polarity, we identified a much larger number of unique annotated metabolites than unique peaks, whereby on average 2.26 chemical formulas were matched to a peak.

**Figure 3.**
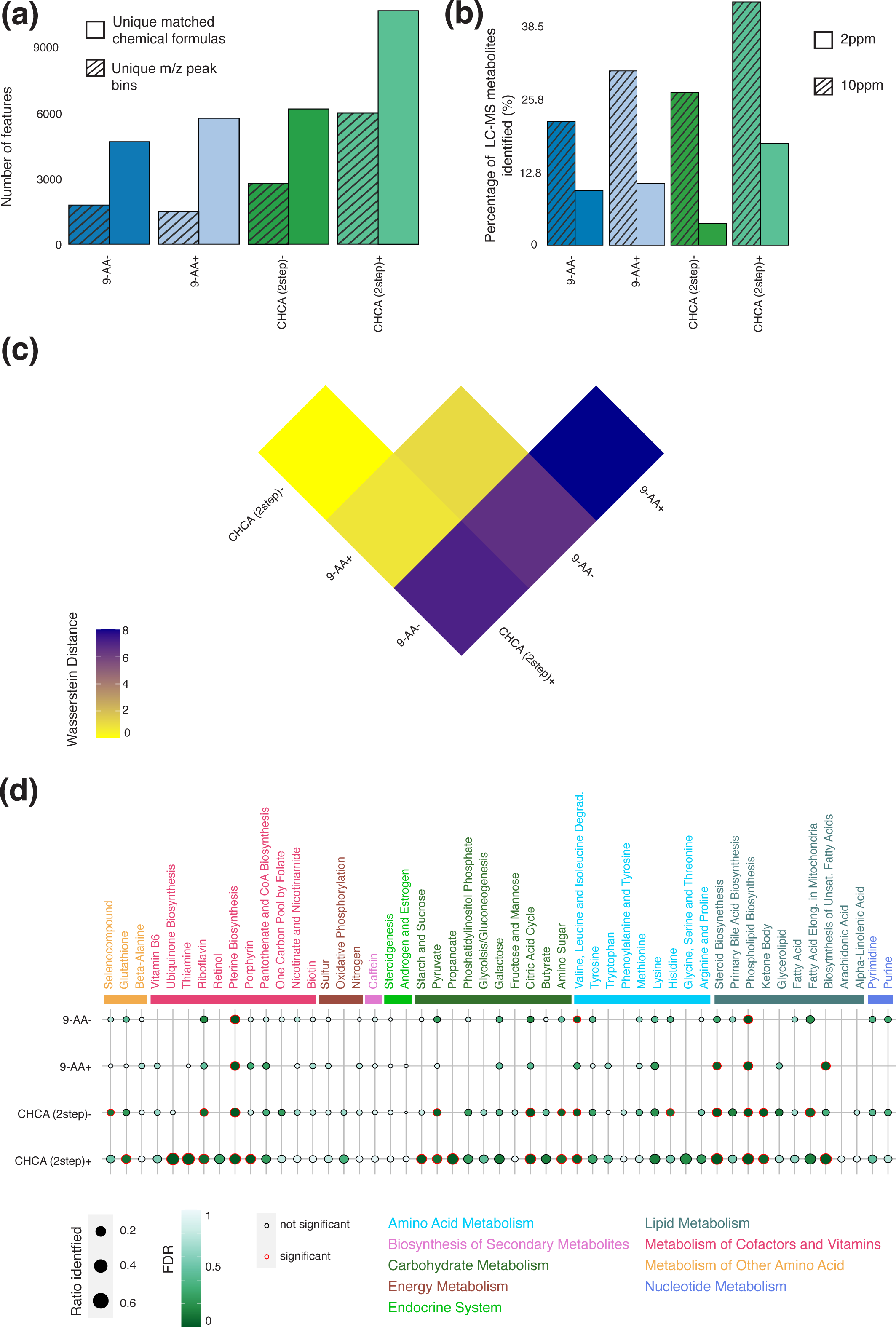
Comparison of different matrix and polarity combinations for detection of lipids in murine glioma. (**a**) Number of unique identified annotated peaks and m/z peaks for each combination; (**b**) Percentage of metabolites detected by LC-MS identified by each combination at 2 and 10ppm thresholds. (**c**) Wasserstein distance between each combination with higher Wasserstein distances between two combinations indicate larger differences between identified peak distribution; (**d**) Ratio of detected metabolites per KEGG pathway and associated FDR for each combination, significantly enriched pathways are indicated with a red outline.

LC-MS on both tumor and normal samples from mouse with CT-2A glioma detected a total of 156 unique annotated lipids. Due to the presence of structural isomers in lipids, the annotation of a m/z value to a specific structure is difficult without fragmentation patterns. To align the LC-MS and MALDI TOF MSI data, we matched peaks detected spatially with the candidate metabolites detected by LC-MS using two different tolerance ranges at 2 and 10 ppm. At 2 ppm, recrystallized CHCA in negative polarity mode had the lowest overlap with less than 10 % (Figure 3b). Maintaining a consistent matching ratio hierarchy, increasing mass tolerance to 10 ppm significantly increased the matching ratio of recrystallized CHCA matrix in negative polarity mode and recrystallized 9-AA in positive polarity mode, possibly due to CHCA’s proton donor and 9-AAs’ proton receiver properties. Hence, using CHCA in negative polarity mode and 9-AA in positive polarity mode might introduce extra noise. Moreover, recrystallized CHCA matrix in positive polarity had the highest number of overlapping peaks at both 2 and 10 ppm owing to the total number of detected peaks.

All matrix and polarity combinations, apart from 9-AA in negative polarity mode, identified similar spectra (Figure 3c). In line with this, 75 % of total number of matched chemical formulas across all matrix and polarity combinations were detected multiple times (Supplementary Material Figure S3c). Owing to the large number of total peaks detected, only recrystallized CHCA matrix in positive polarity mode found more than 23 % of peaks exclusively.

A pathway analysis was conducted to assess whether the different matrix and polarity combinations converged on the detection of similar biological programs. Very similar KEGG pathways were detected across all tested matrix and polarity combinations (Figure 3d). These pathways were largely the same as detected by the small metabolite screen owing to the involvement of both lipids and small molecules in most pathways. As expected, the lipid screen did detect more pathways classified as lipid metabolism and bias towards the detection of such pathways. Pathway enrichment across multiple pathway databases indicated that only recrystallized CHCA matrix in positive polarity mode was biased towards detecting metabolic pathways related to ERBB Family Signaling, which is a key driver pathway in cancer [24] (Supplementary Material Figure S3d).

### 3.3. Differential abundance between tumor and normal regions in murine CT-2A glioma

We next investigated the metabolic differences between healthy and tumor regions in the brain of mice with transplanted CT-2A glioma. Overall, we identified the largest differences between the regions using the recrystallized CHCA matrix in positive polarity mode for both small molecules and lipids (Figures 4a and 4b). Despite detecting many peaks across both ranges, the 9-AA matrix in negative polarity mode identified considerably less differences between the regions than recrystallized CHCA matrix in positive polarity mode. Interestingly, the largest contribution to the differences detected in the recrystallized CHCA matrix in positive polarity mode for small molecules was spermidine, contributing more than 4 % to the overall Wasserstein distance (Supplementary Material Figures S4a). Spermidine has the capacity to improve glucose and lipid metabolism [25], contributing to the promotion of tumor growth. For lipids, no single peak contributed more than 1% to the overall Wasserstein distance (Supplementary Material Figures S4b).

**Figure 4.**
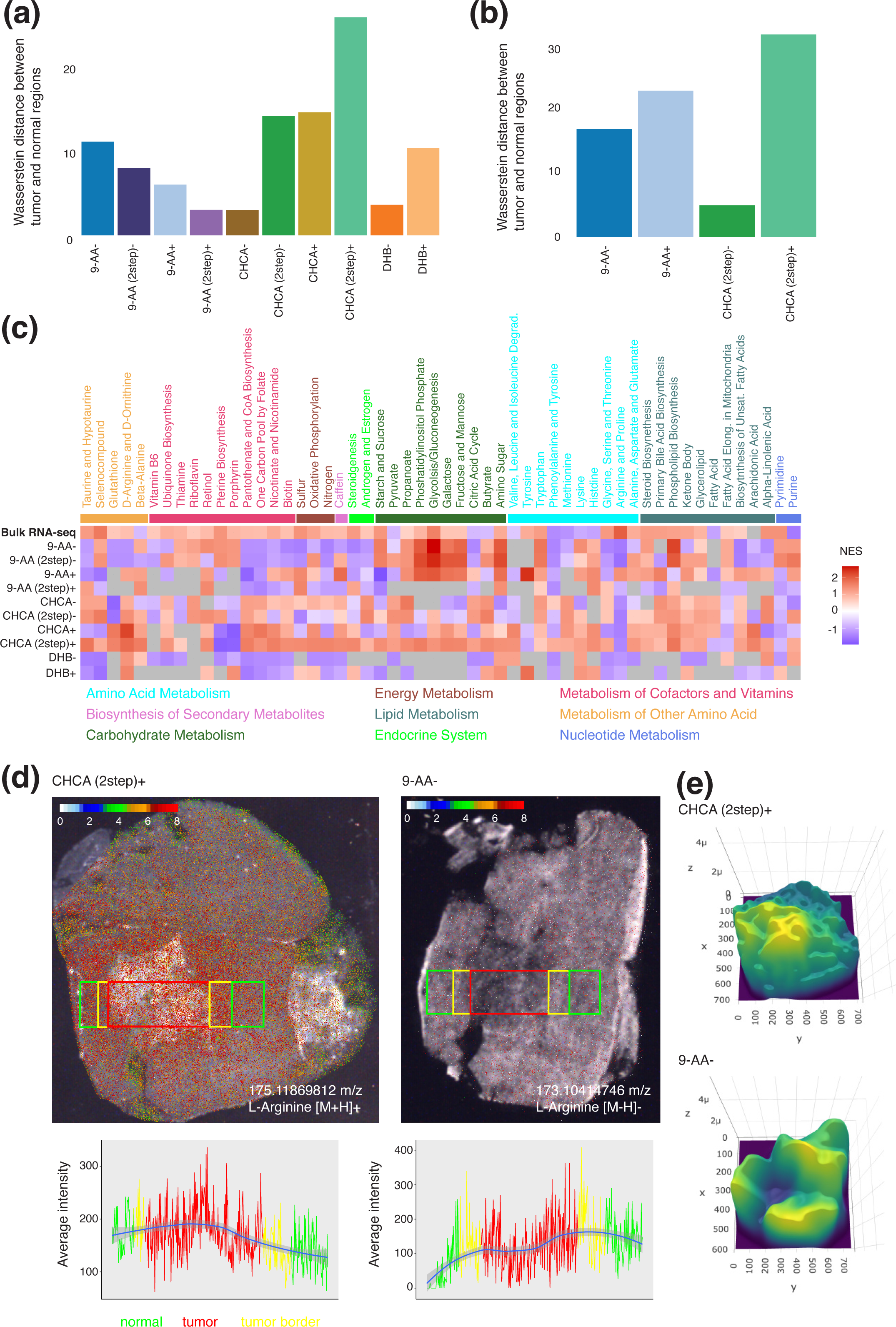
Altered metabolism between tumor and normal regions in murine glioma. (**a**) Wasserstein distance between m/z peak distributions of tumor and normal regions for each combination in small molecule screen; (**b**) Wasserstein distance between m/z peak distributions of tumor and normal regions for each combination in lipid screen; (**c**) Normalized enrichment score for KEGG pathways for each combination and bulk RNA-seq data of tumor and normal regions for small molecule screen; (**d**) Abundance of L-Arginine across tumor (red rectangle), border (yellow rectangle) and adjacent normal (green rectangle) as detected by recrystallized CHCA in positive polarity mode on the left and 9-AA in negative polarity mode on the right, underneath line plot shows the average intensity across the different areas; (**e**) 3D plot of the abundance of L-Arginine for recrystallized CHCA matrix in positive polarity mode (top) and 9-AA matrix in negative polarity mode (bottom).

For the small molecules, we compared the direction of the differential abundance for each matrix and polarity combination (Figure 2b). Of the metabolites identified, 54 % shared the same direction for the differential abundance between tumor and normal in both recrystallized CHCA matrix in positive polarity mode and LC-MS. For all other matrix and polarity combinations, this was below 25 %. At closer inspection, the recrystallized CHCA matrix in positive polarity mode agreed with LC-MS for both reduced and increased metabolites (Supplementary Material Figures S4c). In contrast, the 9-AA matrix in negative polarity mode barely identified any of the same down-regulated metabolites as LC-MS (Supplementary Material Figures S4d).

To further validate our findings, we also conducted RNA-seq on normal and tumor regions from a brain with transplanted CT-2A glioma. To compare the results from RNA-seq to those of MSI, we calculated the normalized enrichment score for KEGG pathways (Figure 4c and Supplementary Material Figures S4e). As for LC-MS, the recrystallized CHCA matrix in positive polarity mode was the most concordant with the RNA-seq results, with 66 % of pathways sharing the direction of overall activity change in the pathway for small molecules and 73 % for lipids. While the 9-AA matrix in negative polarity mode displayed agreement with the LC-MS results, there was less stringency with the RNA-seq results, with 26 % pathways displaying same overall direction in small metabolites and 30 % pathways in lipids. Particularly, all KEGG pathways classified as energy metabolism and endocrine system showed opposite direction changes between tumor and normal.

Examining the distribution of single metabolites across the tissue revealed stark contrasts between the 9-AA in negative polarity mode and the recrystallized CHCA matrix in positive polarity mode. L-Arginine, a conditionally essential amino acid, and required by cancer cells for rapid growth, would be expected to be increased in abundance in the tumor regions. Strikingly, L-Arginine was only detected at high levels using recrystallized CHCA matrix in positive polarity mode and was infrequently detected with 9-AA matrix in negative polarity mode (Figure 4d). Measuring the abundance of L-Arginine spatially across the brain revealed the expected increase in abundance of the amino acid specifically in the tumor region (Figure 4e), while 9-AA in negative polarity mode was significantly diminished across the tumor. Thus, recrystallized CHCA matrix in positive polarity mode appears to be most suited to the normal and tumor brain microenvironment.

### 3.4. Matrix selection for metabolite detection in human IDH-mutant glioma

To test whether our findings were transferable across species and brain tumor subtype, we applied the two best performing matrix and polarity combinations on resected tumor sample from a patient with *IDH*-mutant glioma. We found similar trends with regards to the number of detected peak bins and annotated chemical formulas as was observed for murine CT-2A glioma for both small metabolites and lipids (Figures 5a and 5b). The recrystallized CHCA matrix in positive polarity mode detected 1.64 times more peaks bins for the small molecules and 2.34 times more for the lipids than the 9-AA matrix in negative polarity mode. This resulted in 227 % more chemical formulas for the small molecules and 160 % more for the lipids detected in the recrystallized CHCA matrix in positive polarity mode compared to the 9-AA matrix in negative polarity mode.

**Figure 5.**
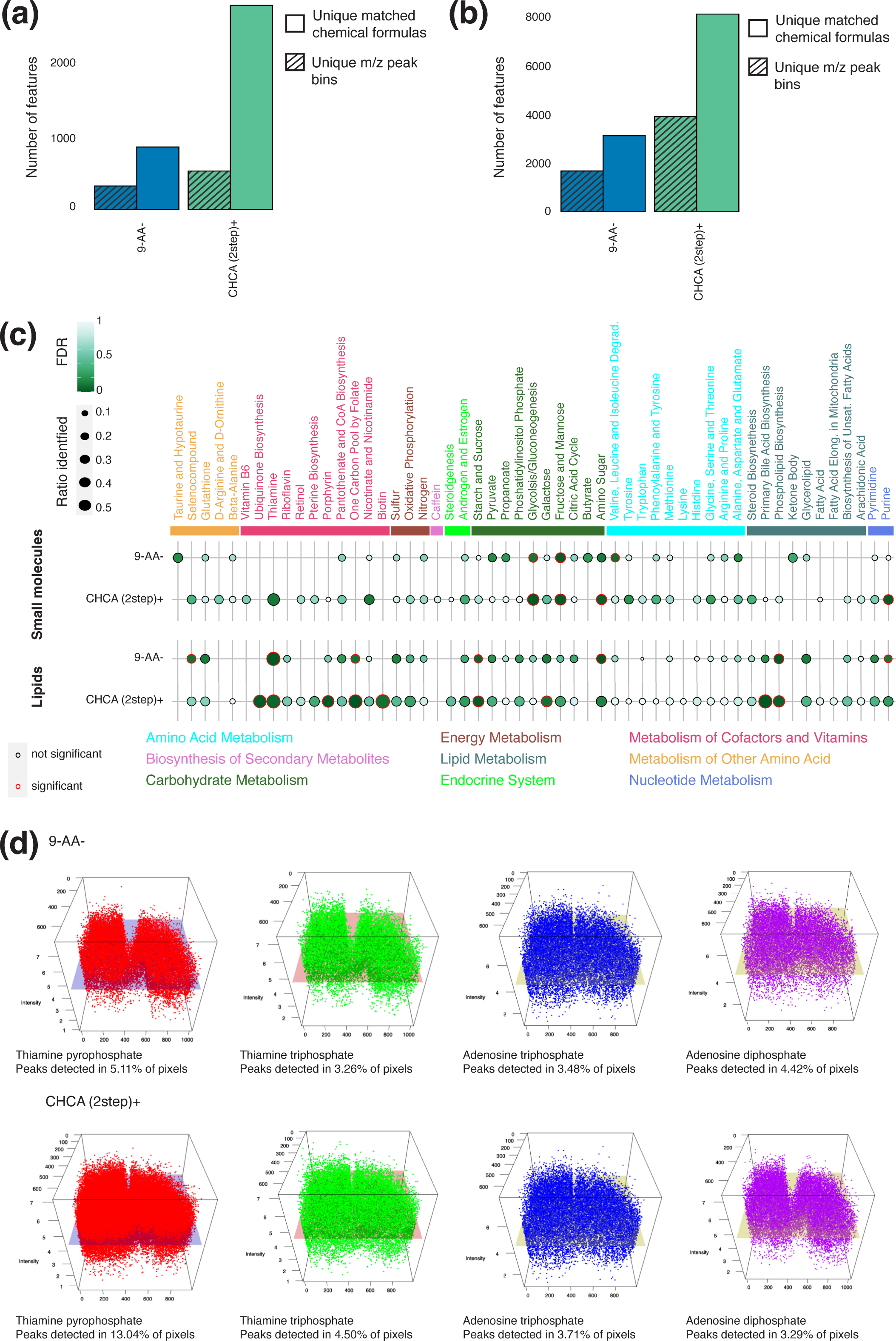
Lipid and small molecules in a tumor from a patient with *IDH1*-mutated glioma. (**a**) Number of unique identified annotated peaks and m/z peak bins for each combination in small molecule screen; (**b**) Number of unique identified annotated peaks and m/z peak bins for each combination in lipid screen; (**c**) Ratio of detected metabolites per KEGG pathway and associated FDR for each combination separated by screen, significantly enriched pathways are indicated with a red outline; (**d**) 3-D visualization of selected metabolites in the Thiamine pathway across the tumor section for 9-AA matrix in negative polarity mode (top) and recrystallized CHCA matrix in positive polarity mode (bottom) in the lipid screen.

As previously, we conducted a pathway analysis for both lipids and small molecules with very similar results in terms of detection (Figure 5c). For both, most pathways represented in the 9-AA matrix in negative polarity mode were represented in the recrystallized CHCA matrix in positive polarity mode except for two lipid pathways and Butyrate in the small molecule screen. However, the 9-AA matrix in negative polarity mode missed several pathways belonging to the group of Metabolism of Cofactors and Vitamins, required for chemical reactions in the cell. In contrast to results observed for murine CT-2A glioma, recrystallized CHCA matrix in positive polarity mode was enriched for metabolites in pathways grouped as Carbohydrate Metabolism rather than Energy Metabolism. This is likely a reflection of the biology of decreased glucose uptake in *IDH*-mutant tumors compared to *IDH*-wildtype tumors, such as CT-2A glioma [26].

Finally, we visualized metabolites of the Thiamine pathway which is statistically significantly enriched in both matrices for lipids (Figure 5d) and observed that for most metabolites enriched in the pathway, the recrystallized CHCA matrix in positive polarity mode were detectable across many more pixels than with the 9-AA matrix in negative polarity mode. Furthermore, the average intensity of the metabolites was higher.

## 4. Discussion

The selection of matrix mixture, deposition technique, and polarity setting have a crucial impact on metabolite detection in MALDI MSI. To guide matrix selection for the study of brain cancers, we tested several combinations of matrices, deposition techniques, and polarity settings across brain cancer types and species for both small molecules and lipids. We found that the use of a recrystallized CHCA matrix in positive polarity mode for MALDI MSI in glioma allowed us to detect the greatest number of metabolites across species and brain cancer types. Comparison with LC-MS data generated from the same samples also showed the greatest overlap for this matrix and polarity combination. The commonly applied 9-AA matrix in negative polarity mode in the glioma setting had inferior performance in terms of the number of metabolites detected and matching with LC-MS data.

The superiority of the recrystallized CHCA matrix in positive polarity mode among the tested combinations was also confirmed when comparing differential abundance between tumor and normal regions across different technologies. In particular, a high degree of concordance was observed between metabolite peaks detected using the CHCA matrix in positive polarity mode and RNA expression derived from RNA-seq data, increasing confidence in the results. Additionally, an example of spatial abundance of L-Arginine, an important metabolite involved in cancer growth, revealed sparsity of detection in the commonly applied 9-AA matrix in negative polarity mode compared to the recrystallized CHCA matrix in positive polarity mode. This improved detection is possibly the result of the peculiar features of ionic liquid matrices leading to smaller spot sizes, minimization of experimental variation, decrease in matrix cluster formation and reduced fragmentation [27].

Beyond guiding matrix selection specifically for the study of metabolic alterations in brain cancer, our work establishes an experimental and computational workflow to determine the optimal matrix for any spatial metabolomics study. This workflow includes ascertaining gross differences between matrices, biological interpretation of differences, and comparison with other techniques such as LC-MS and RNA-seq. In particular, the comparison with other techniques allows assessment of the overlap between results with a greater overlap likely representing a greater number of real observed differences, thus providing an objective quality measure. Application of our workflow will enable objective matrix selection and improve MALDI-TOF MSI studies in the future.

## Supplementary Materials

The following supporting information can be downloaded at: www.mdpi.com/xxx/s1, Figure S1: Validation of MALDI-TOF using bulk RNA-seq; Figure S2: Further comparisons between matrix and polarity combinations for MALDI-TOF for small molecules; Figure S3: Further comparisons between matrix and polarity combinations for MALDI-TOF for lipids; Figure S4: Extended differential metabolite abundance between tumor and normal regions.

## Author Contributions

Conceptualization, S.F, S.A.B., and J.R.W.; methodology, S.F., L.F., T.L., M.J.M., and V.N.; sample preparation, A. V., S.J.O., Z.M. and S.A.B.; software, L.F. and T.L.; formal analysis, T.L.; resources, S.F, S.A.B., M.J.M., and J.R.W.; data curation, V.N., A.V., B.N., A.P., and D.S.; writing—original draft preparation, S.F., T.L., and S.A.B.; writing—review and editing, S.F., S.A.B., L.F., T.L., V.N., M.J.M., A.V., A.P., B.N., D.S. and J.R.W.; visualization, S.F. and T.L.; supervision, S.F. and S.A.B.; project administration, S.F. and S.A.B.; funding acquisition, S.F, S.A.B., M.J.M., and J.R.W. All authors have read and agreed to the published version of the manuscript.

## Institutional Review Board Statement

The study was conducted in accordance with the Declaration of Helsinki and approved by the Melbourne Health Ethics Committee (protocol code 2020.214) and the Walter and Eliza Hall Human Research Ethics Committee (protocol code 21/21 approved 28/6/2021). The animal study protocol was approved by the Ethics Committee of the Walter and Eliza Hall Institute of Medical Research (protocol code 2022.004 approved 22/03/2022).

## Informed Consent Statement

Informed consent was obtained from all subjects involved in the study.

## Data Availability Statement

Data is available during the review period on request to the author or editor. All code is available via https://github.com/BCRL-tylu/Matrix-optimisation.

## Acknowledgments

We thank S. Stylli, K. Drummond, and J. Dimou for expert curation of the Royal Melbourne Hospital Neurosurgery Brain and Spine Tumor Tissue Bank, M. Bisignano for assistance at the RMH Anatomical Pathology Department and E. Tsui and A. Maluenda for assistance at the WEHI Histology core.

