## Supplementary Materials for "Matrix selection for the visualization of small molecules and lipids in brain tumors using untargeted MALDI-TOF mass spectrometry imaging"

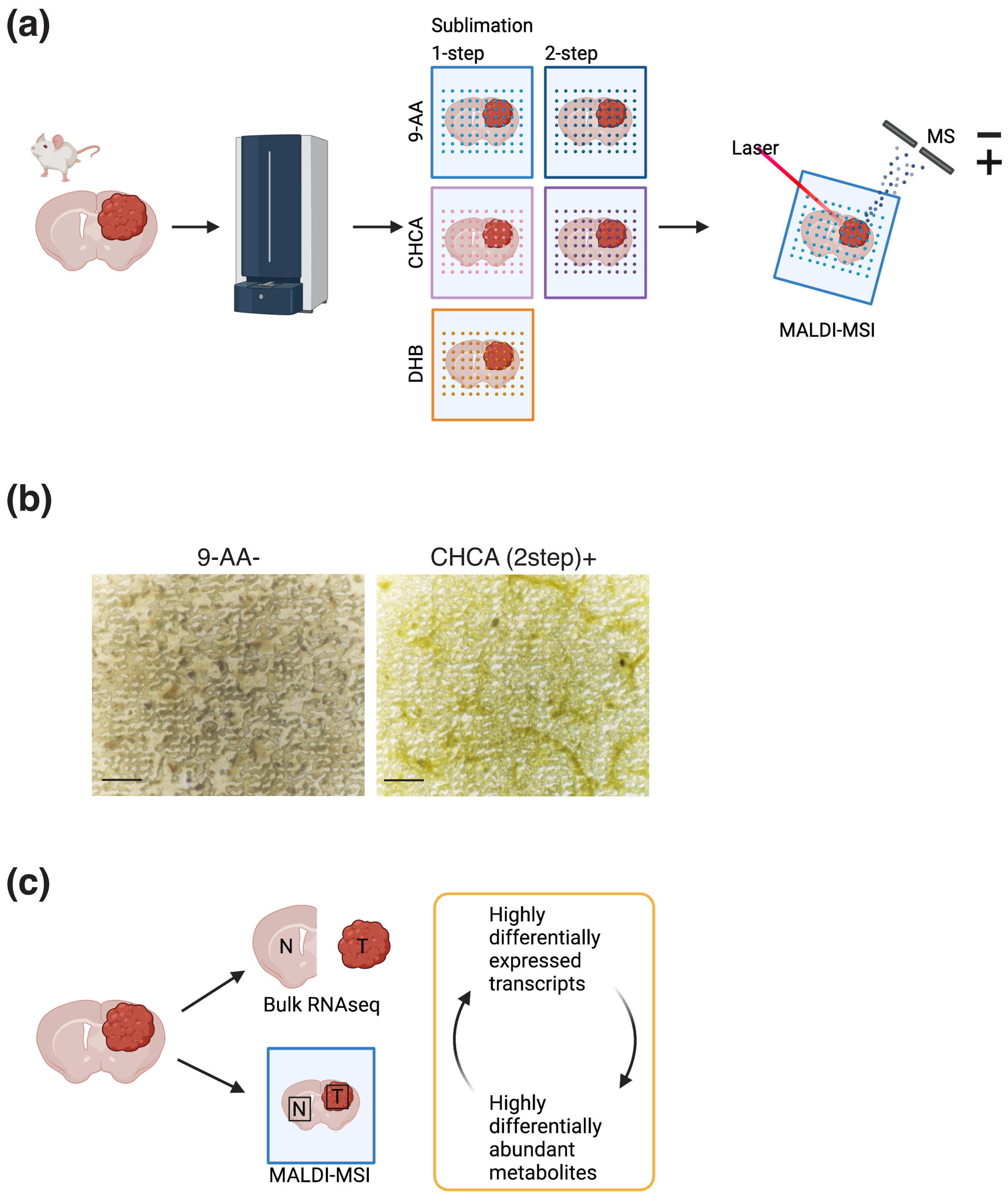
(a)
Sublimation
1-step
2-step
9-AA
CHCA
DHB
Laser
MS
MALDI-MSI
(b)
9-AA-
CHCA (2step)+
(c)
N
T
Bulk RNAseq
N
T
MALDI-MSI
Highly differentially expressed transcripts
Highly differentially abundant metabolites

**Supplementary Material Figure S1.** Experimental designs. (**a**) Experimental design for matrix comparison with CT-2A tumors; (b) Microscopy of MALDI-analyzed CT-2A tumors highlighting laser destruction of tissue; scale, 50 μm; (**c**) Experimental design for validation of MALDI-TOF using bulk RNA-seq.

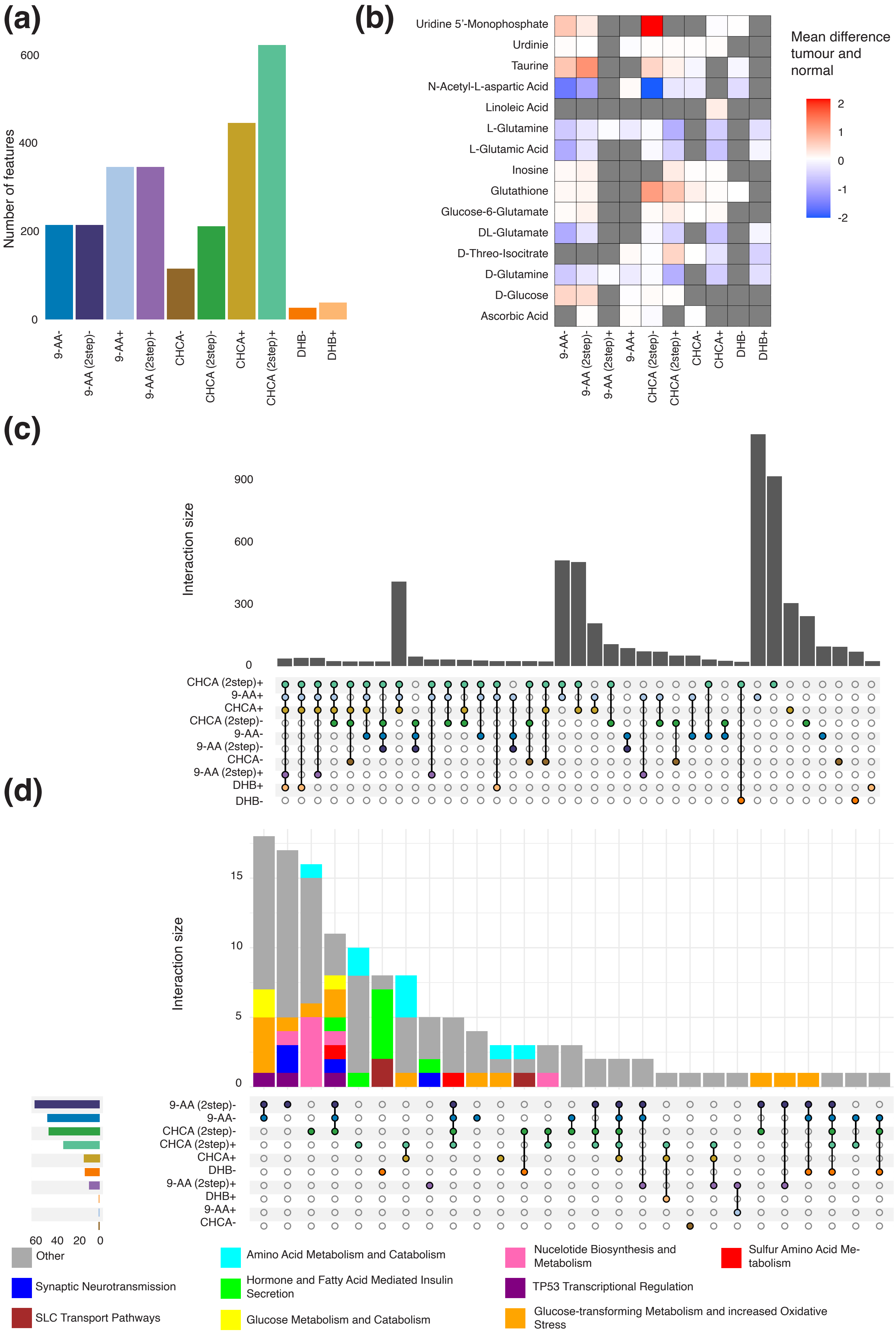
(a)
Number of features
0
200
400
600
9-AA-
9-AA (2step)-
9-AA+
9-AA (2step)+
CHCA-
CHCA (2step)-
CHCA+
CHCA (2step)+
DHB-
DHB+
(b)
Uridine 5'-Monophosphate
Urdinie
Taurine
N-Acetyl-L-aspartic Acid
Linoleic Acid
L-Glutamine
L-Glutamic Acid
Inosine
Glutathione
Glucose-6-Glutamate
DL-Glutamate
D-Threo-Isocitrate
D-Glutamine
D-Glucose
Ascorbic Acid
9-AA-
9-AA (2step)-
9-AA (2step)+
9-AA+
CHCA (2step)-
CHCA (2step)+
CHCA-
CHCA+
DHB-
DHB+
Mean difference tumour and normal
2
1
0
-1
-2
(c)
Interaction size
0
300
600
900
CHCA (2step)+
9-AA+
CHCA+
CHCA (2step)-
9-AA-
9-AA (2step)-
CHCA-
9-AA (2step)+
DHB+
DHB-
(d)
Interaction size
0
5
10
15
9-AA (2step)-
9-AA-
CHCA (2step)-
CHCA (2step)+
CHCA+
DHB-
9-AA (2step)+
DHB+
9-AA+
CHCA-
60
40
20
0
Other
Amino Acid Metabolism and Catabolism
Nucelotide Biosynthesis and Metabolism
Sulfur Amino Acid Metabolism
Synaptic Neurotransmission
Hormone and Fatty Acid Mediated Insulin Secretion
TP53 Transcriptional Regulation
SLC Transport Pathways
Glucose Metabolism and Catabolism
Glucose-transforming Metabolism and increased Oxidative Stress

**Supplementary Material Figure S2.** Further comparisons between matrix and polarity combinations for MALDI-TOF for small molecules. (**a**) Number of unique background peaks for each combination in small molecule screen; (**b**) Mean difference in abundance of metabolites of interest between tumor and normal regions with grey color representing metabolites not detectable; (**c**) Upset plot representing a Venn diagram of sizes of overlaps between detectable chemical formulas for different sets of matrix and polarity combinations; (**d**) Upset plot representing a Venn diagram of sizes of overlaps between enriched pathways for different sets of matrix and polarity combinations colored by semantic groups of pathways.

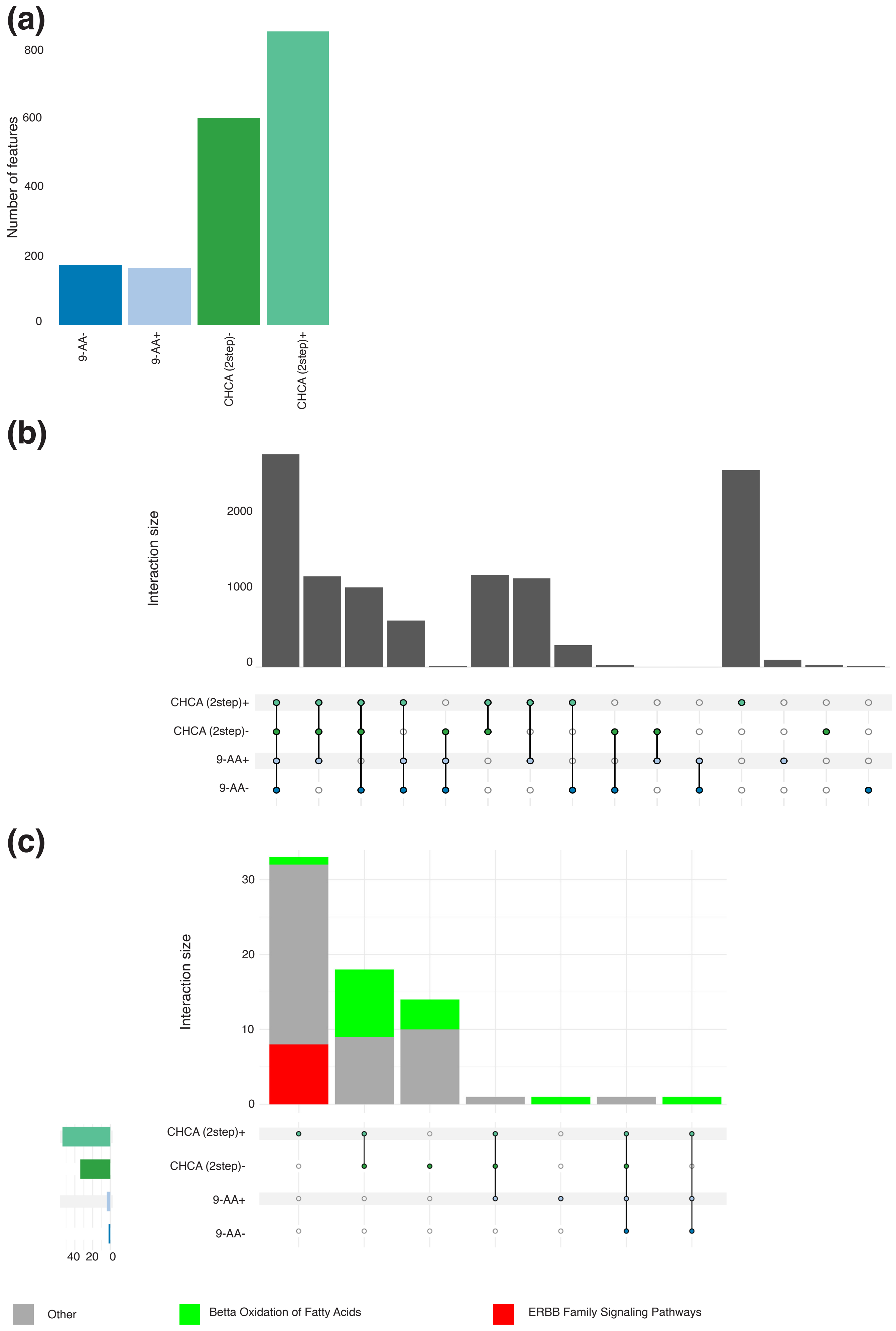

(a)
Number of features
800
600
400
200
0
9-AA-
9-AA+
CHCA (2step)-
CHCA (2step)+
(b)
Interaction size
2000
1000
0
CHCA (2step)+
CHCA (2step)-
9-AA+
9-AA-
(c)
Interaction size
30
20
10
0
CHCA (2step)+
CHCA (2step)-
9-AA+
9-AA-
40 20 0
Other
Betta Oxidation of Fatty Acids
ERBB Family Signaling Pathways

**Supplementary Material Figure S3.** Further comparisons between matrix and polarity combinations for MALDI-TOF for lipids. (**a**) Number of unique background peaks for each combination in small molecule screen; (**b**) Upset plot representing a Venn diagram of sizes of overlaps between detectable chemical formulas for different sets of matrix and polarity combinations; (**c**) Upset plot representing a Venn diagram of sizes of overlaps between enriched pathways for different sets of matrix and polarity combinations colored by semantic groups of pathways.

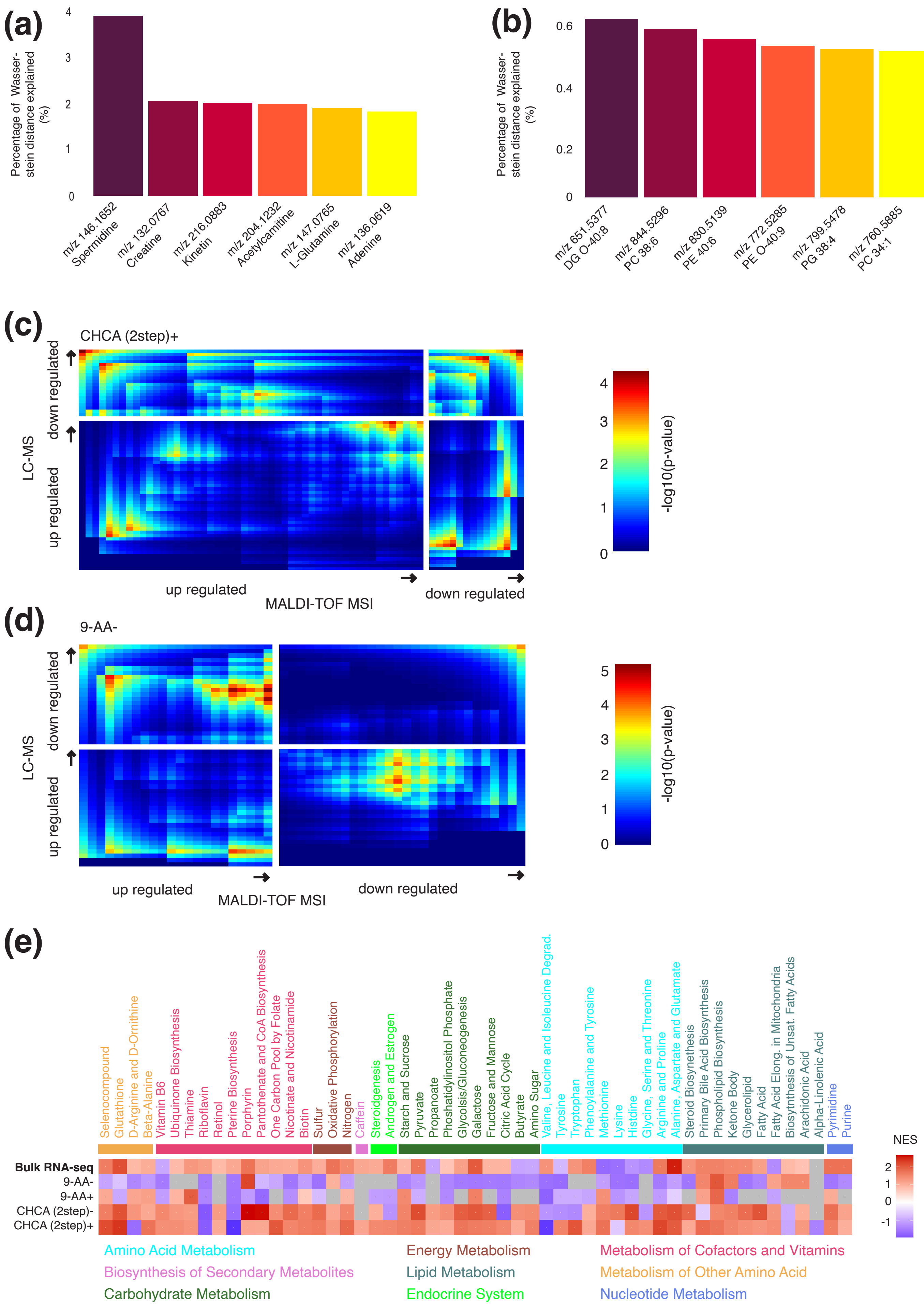
(a)
Percentage of Wasser-stein distance explained (%)
m/z 146.1652 Spermidine
m/z 132.0767 Creatine
m/z 216.0883 Kinetin
m/z 204.1232 Acetylcarnitine
m/z 147.0765 L-Glutamine
m/z 136.0619 Adenine
(b)
m/z 651.5377 DG O-40:8
m/z 844.5296 PC 38:6
m/z 830.5139 PE 40:6
m/z 772.5285 PE O-40:9
m/z 799.5478 PG 38:4
m/z 760.5885 PC 34:1
(c)
CHCA (2step)+
LC-MS
down regulated
up regulated
MALDI-TOF MSI
-log10(p-value)
(d)
9-AA-
(e)
Selenocompound
Glutathione
D-Arginine and D-Ornithine
Beta-Alanine
Vitamin B6
Ubiquinone Biosynthesis
Thiamine
Riboflavin
Retinol
Pterine Biosynthesis
Porphyrin
Pantothenate and CoA Biosynthesis
One Carbon Pool by Folate
Nicotinate and Nicotinamide
Biotin
Sulfur
Oxidative Phosphorylation
Nitrogen
Caffein
Steroidgenesis
Androgen and Estrogen
Starch and Sucrose
Pyruvate
Propanoate
Phoshatidylinositol Phosphate
Glycolsis/Gluconeogenesis
Galactose
Fructose and Mannose
Citric Acid Cycle
Butyrate
Amino Sugar
Valine, Leucine and Isoleucine Degrad.
Tyrosine
Tryptophan
Phenoylalanine and Tyrosine
Methionine
Lysine
Histdine
Glycine, Serine and Threonine
Arginine and Proline
Alanine, Aspartate and Glutamate
Steroid Biosynthesis
Primary Bile Acid Biosynthesis
Phospholipid Biosynthesis
Ketone Body
Glycerolipid
Fatty Acid
Fatty Acid Elong. in Mitochondria
Biosyntnthesis of Unsat. Fatty Acids
Arachidonic Acid
Alpha-Linolenic Acid
Pyrimidine
Purine
Bulk RNA-seq
9-AA-
9-AA+
CHCA (2step)-
CHCA (2step)+
NES
Amino Acid Metabolism
Energy Metabolism
Metabolism of Cofactors and Vitamins
Biosynthesis of Secondary Metabolites
Lipid Metabolism
Metabolism of Other Amino Acid
Carbohydrate Metabolism
Endocrine System
Nucleotide Metabolism

**Supplementary Material Figure S4.** Extended differential metabolite abundance between tumor and normal regions. (**a**) Percentage of total Wasserstein distance between tumor and normal regions explained by top 6 differentially abundant m/z peak bins for small molecule screen; (**b**) Percentage of total Wasserstein distance between tumor and normal regions explained by top 6 differentially abundant m/z peak bins for lipid screen; (**c**) Rank-rank hypergeometric overlap plots between LC-MS and MALDI-TOF MSI differential abundance results of small molecules for recrystallized CHCA matrix in positive polarity mode; smaller p-values indicate larger overlap between ranked lists; (**d**) Rank-rank hypergeometric overlap plots between LC-MS and MALDI-TOF MSI differential abundance results of small molecules for 9-AA matrix in negative polarity mode; smaller p-values indicate larger overlap between ranked lists; (**e**) Normalized enrichment score for KEGG pathways for each combination and bulk RNA-sequencing of tumor and normal regions for lipid screen.
